# celltypeEnrich: a consensus-based scRNA-seq cluster annotation tool

**DOI:** 10.64898/2026.09.15.751735

**Authors:** S. Rutledge, G. Tuteja

## Abstract

**Motivation:** Single-cell RNA sequencing (scRNA-seq) cluster annotation is a critical step in data analysis. Current methods are time-consuming, difficult to reproduce, or limited in tissue or species coverage.

**Results:** We developed celltypeEnrich, a cluster-level annotation tool that uses a hypergeometric test to identify enrichment of cell-type-specific genes from input gene lists. Enrichment results from up to 26 reference datasets are used to determine a consensus annotation. Benchmarking using scRNA-seq datasets from three tissues spanning two species showed 62-72% annotation accuracy for celltypeEnrich, generally outperforming other tools, which had either lower accuracy, incomplete tissue coverage, or the need for parameter optimization. The performance of celltypeEnrich remained stable when input gene lists were down-sampled to 25% of their original size.

**Availability and Implementation:** celltypeEnrich is freely available at (https://celltypeenrich.gdcb.iastate.edu) as an R Shiny web application under the MIT license for non-profit academic use.

**Contact:** For further questions or assistance, please contact Dr. Tuteja at.

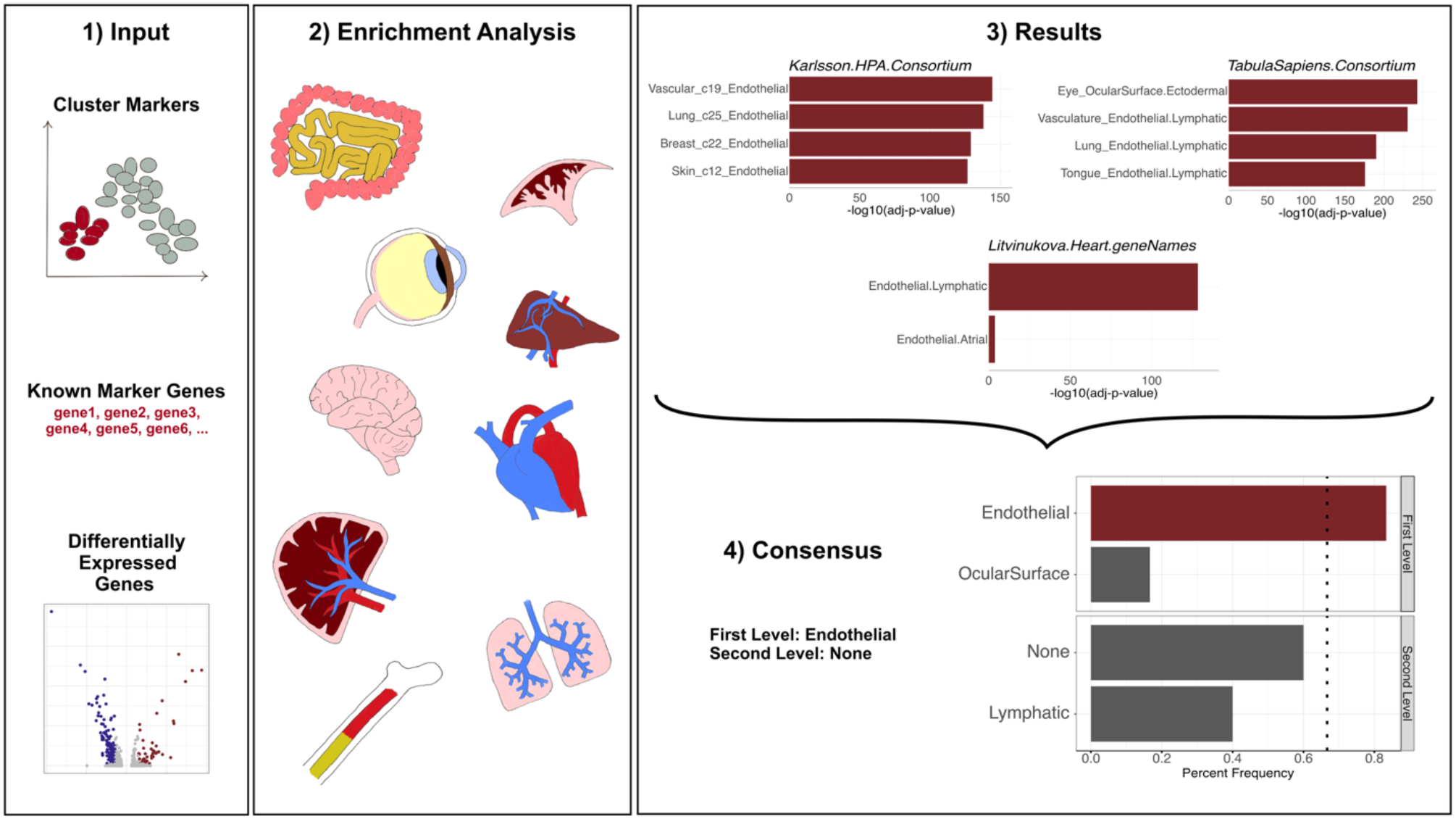

## 1) Introduction

Since its introduction in 2009^1^, single-cell RNA sequencing (scRNA-seq) has allowed researchers to profile transcriptomic heterogeneity at the level of individual cells. Advances in scRNA-seq technology have reduced costs and increased both the number of cells profiled and the depth of transcript coverage per cell, contributing to its adoption in diverse biological contexts. The analysis pipeline for scRNA-seq data has remained relatively consistent and includes preprocessing and dimensional reduction to obtain groups, or clusters, of cells with similar transcriptional profiles. These clusters must then be reliably annotated with their cellular identity for downstream analyses, such as trajectory and cell-cell communication analyses. However, cluster annotation remains a major challenge^2^.

A common approach to cell-type annotation involves researchers visualizing well-known marker genes on dimensional-reduction plots (e.g. UMAP or tSNE), often relying on only one or two marker genes per cell type. This largely manual approach can be misleading, as clusters often express marker genes from multiple cell types, which may not be jointly considered during annotation. These issues can lead to inaccurate annotations and ultimately impact downstream analyses.

Considering these limitations with manual annotation, numerous computational tools have been developed for scRNA-seq cluster- or cell-level annotation. Many annotation tools rely on reference databases^3^ or pretrained models^4–6^, assuming that the data being annotated resembles the references. This assumption does not apply when the reference resources have limited tissue coverage or are restricted to a narrow range of model organisms. Even for an individual tissue, using multiple reference datasets or models often produces inconsistent annotations for the same cell or cluster.^7^ Several tools leverage large consortium-generated atlases^6,8–10^, such as the Human Cell Atlas^11^, Human Protein Atlas^12^ (HPA), and Tabula Sapiens^13^, which generate and compile expression data from multiple tissues into a single reference dataset. While these consortium references include a wide array of cell types, highly specific cell types or subtypes are often missing. Other tools^14–16^ depend on user-provided information, such as curated marker lists, which can be problematic when these lists are incomplete or not well established for certain cell types. In addition, many existing annotation tools^3–5^ accept only human or mouse genes as input, limiting their usability for researchers working with other model organisms who lack bioinformatics expertise.

To address these challenges, we developed celltypeEnrich, a cluster-level annotation tool that tests whether an input gene list is significantly enriched for cell-type-specific genes calculated from multiple independent datasets. By analyzing enrichment results across these datasets, celltypeEnrich determines a consensus annotation, where recurrent significant enrichments provide greater confidence for the assigned cell type. celltypeEnrich provides consistent and high-confidence annotations across tissues and supports analysis of multiple model organisms.

## 2) Methods

### 2.1) scRNA-seq datasets used in celltypeEnrich

#### 2.1.1) Collecting scRNA-seq Datasets

scRNA-seq datasets^12,13,17–35^ from each major human organ system were obtained from either the Teichmann group atlas websites (https://www.teichlab.org/data)^19,21,22,25,27,31^ or through the CZ cellxgene website^9^ (Supplemental Table 1). Data from disease samples were excluded. Multiple RDS Seurat-object files were available and downloaded for the Yan Retina^34^, Elmentaite Gut^17^, and Vento-Tormo Placenta^30^ datasets, and all of the data for each individual tissue were integrated into a single dataset using the standard Seurat^36^ pipeline. The Kernfeld Thymus^23^ dataset had multiple raw data text files which were merged prior to processing with the standard Seurat pipeline. In addition to obtaining scRNA-seq datasets from individual tissues, we downloaded processed data for 30 tissues from the Human Protein Atlas^12^ and 23 tissues from Tabula Sapiens^13^.

#### 2.1.2) Normalizing scRNA-seq Datasets

For each individual tissue dataset, we compared the variance in the downloaded data files to both log-normalized and CLR-cell-normalized data using variance plots^37^. The normalization method yielding the most stable variance was used for downstream analyses. For the four datasets that needed to be integrated prior to analysis in 2.1.1, we compared variance plots between log-normalization and CLR-cell-normalization. The normalization method used for each dataset is provided in Supplemental Table 1. The two consortium datasets included the integrated data from all assayed tissues in a single file and were used as provided for the downstream analyses.

#### 2.1.3) Cell type relabeling

Cell types were relabeled to ensure consistency within and across datasets. Each label was re-organized into a hierarchical pattern separated by periods, where the first word/hierarchical level represents the broadest cell category and the last word/hierarchical level represents the most specific cell category, often reflecting a region of origin or a defining gene with which the cell type was categorized (for example, Epithelial.OvarianSurface and Endothelial.SEMA3G). For the two consortium datasets where several different tissues are part of one large dataset, there is a prefix naming the tissue of origin. For the Human Protein Atlas^12^ dataset, the cluster number is included in the prefix since there were multiple distinct clusters labeled with the same cell type within each tissue. Multi-word cell-type names were concatenated into a single word (e.g. ‘Natural Killer’ was relabeled to ‘NaturalKiller’). The full list of cell types included in celltypeEnrich is in Supplemental Table 2.

### 2.2) Determining cell-type-specific genes for each reference dataset

celltypeEnrich evaluates whether a given gene list is overrepresented for cell-type-specific genes using gene sets derived from each reference dataset. To optimally generate these cell-type-specific gene sets, we considered both the average expression of each gene within a cell type and its relative expression compared to all other cell types, as described in detail below. For each cell type in each reference dataset, the average expression profile was obtained with the AverageExpression function in Seurat^36^. Then the average expression data-frame was used as input for the teGeneRetrieval function in TissueEnrich^38^, which uses HPA’s algorithm^39^ to determine if a gene is cell-type-specific. HPA’s algorithm defines cell-type-specific genes based on (1) fold-change in expression between a cell type of interest and all other cell types, and (2) an expression threshold where a gene must be expressed above a pre-defined threshold. These metrics have been widely used to define tissue- and cell-type-specific genes and have formed the basis of several gene enrichment algorithms and software packages^38–44^.

The fold-change threshold was initially set to the default value of 5 and subsequently lowered to 3 to test a range of thresholds. To determine an expression threshold for each reference dataset, the distribution of non-zero expression values was calculated, and expression thresholds were defined using the 50th and 75th percentiles of this distribution. Together, these fold-change and expression thresholds yielded multiple parameter combinations, each producing a corresponding set of cell-type-specific genes for every cell type in each reference dataset.

The HPA algorithm^39^ defines a gene as ‘Group-Enriched’ if it is highly expressed in 2-7 cell types. Given the large number of repeated cell types (i.e. the same cell type across several different tissues) present in each of the two consortium datasets, the parameter defining how many cell types to use as a group for the ‘Group-Enriched’ category of cell-type-specific genes was tested at 15 and 30, in addition to the default of 7, for these datasets. For the Van Zyl Lens and Cornea datasets, the total number of cell types present in the datasets was less than the default for the ‘Group-Enriched’ parameter, so the parameter was lowered to 3 and 5, respectively. In all other cases, this parameter was kept at the default.

To then select the optimal parameter combination for each reference dataset, an independent scRNA-seq dataset was identified for each tissue (Supplemental Table 3, ‘Marker List Source’ Column), for which marker gene lists were provided in the original publication. For each independent marker list, enrichment analysis was performed across all parameter combinations using the teEnrichmentCustom function from TissueEnrich^38^, which applies a hypergeometric test to assess enrichment for cell-type-specific genes. Only enrichments with an adjusted p-value < 0.05 were retained. Then, enrichment results were categorized as ‘Correct,’ when all of the top 10% of enriched cell types matched the annotated cell type reported in the original publication; ‘Mixed,’ when the top 10% had both correct and incorrect cell-type enrichments; or ‘Incorrect,’ when none of the top 10% of enrichments corresponded to the cell type originally reported. If a marker list had no significant enrichments, it was categorized as ‘No-Enrichments.’ The parameter combination with the most ‘Correct’ evaluations was generally used to define the cell-type-specific genes for the reference scRNA-seq dataset that was being evaluated (Supplemental Table 3, ‘expressedGeneThreshold’ and ‘foldChangeThreshold’ Columns). If the number of evaluations per category was consistent across the parameter combinations, then the parameter combination with the more stringent thresholds was selected.

### 2.3) Defining a Consensus Annotation After Enrichment Analysis

Following enrichment analysis for an input gene list, celltypeEnrich derives a consensus annotation to summarize annotation results across the reference datasets. For each reference dataset, cell-type annotations corresponding to enrichments within 10% of the highest negative log10-adjusted p-value (the top-most enriched cell types) are retained. These annotations are then tabulated by frequency at each hierarchical annotation level (see section 2.1.3), starting from the broadest level. A consensus annotation is assigned when a given cell-type label occurs in at least two-thirds of the top-most enriched annotations at a given level. If a consensus is identified, enrichments that do not match the consensus label are removed before proceeding to consensus evaluation at the next, more specific, annotation level. This process continues iteratively through the annotation hierarchy. The tabulated frequencies are provided as a plot in the R shiny application, whether or not a consensus is identified, for each level analyzed.

### 2.4) Parameter Sweeps for Tools Used in the Benchmark Analysis

Current annotation tools can be broadly grouped into those that annotate clusters or individual cells (Supplemental table 4). Since celltypeEnrich annotates clusters using user-provided marker gene lists, we first focused on cluster-level annotation tools that use the same type of input. Of those identified (Supplemental table 4), only EasyCellType^3^ and WebCSEA^6^ accepted the same input format and had been updated within the last three years. To further compare celltypeEnrich with methods that use gene expression matrices as input, we included SingleR^4^, a widely used annotation tool that has been evaluated in multiple benchmarking studies and has been shown to perform comparably to other recent tools^3,45–58^.

EasyCellType^3^ has two options for which statistical test to use, the Fisher’s Exact Test and the GSEA test, and three options for the cell-type marker database to use as a reference, CellMarker^59^, Clustermole^60^, and Panglao^61^. Therefore, EasyCellType has six parameter combinations: the Fisher’s Exact Test with the CellMarker, Panglao, or Clustermole database and the GSEA test with the CellMarker, Panglao, or Clustermole database. For all six combinations, we first labeled results without any significant results as ‘NoResults’. Then each set of significant results was filtered to the top 10% of enrichments, and these were classified as ‘Correct’, ‘Incorrect’, and ‘Mixed’, as described in section 2.2.

SingleR^4^ uses predefined reference datasets to label either clusters or cells. When benchmarking with SingleR, we tested both of the bulk RNA-seq reference datasets (Human Primary Cell Atlas^62^ and the Blueprint Encode datasets^63,64^), which include multiple tissues, and scRNA-seq models that were appropriate for the dataset being tested: the Zillonis Lung Tumor dataset and the He Organ Atlas dataset. The He Organ Atlas dataset had several annotations per cell, which were tested separately. Any cell with no assigned labels in the models was removed prior to testing. For all iterations of scRNA-seq reference dataset testing, the reference dataset’s gene expression matrix was log-normalized with the logNormCounts function. Low confidence cell-type assignments (labeled as NA) were considered similar to non-significant results and were classified as ‘NoResults.’ Otherwise, the cluster assignments were classified as ‘Correct’ or ‘Incorrect’ based on the expected cell type.

For WebCSEA^6^, marker gene lists were submitted through the web interface.

## 3) Results

### 3.1) Implementation and Usage of celltypeEnrich

celltypeEnrich uses a hypergeometric test to assess enrichment for cell-type-specific genes and is implemented as an interactive R Shiny web application. The website is organized into four pages. The ‘Home’ page has general information about the tool and links to the other three pages. The ‘Getting Started’ page has a step-by-step tutorial on using the tool, while the ‘Frequently Asked Questions’ page addresses common user queries. The ‘Cell-type Gene Enrichment’ page hosts the R Shiny application, where users can analyze their gene list to assess enrichment for cell-type-specific genes. Each parameter for the web application is described below.

#### 3.1.1) Input Options

celltypeEnrich requires three inputs (Figure 1). First, the user must specify the species of origin for their input gene list (Figure 1A). While the tool uses human datasets for the enrichment analysis, the user may input a gene list from seven additional species: mouse, rat, pig, sheep, zebrafish, crab-eating macaque, and macaque. If a non-human species is selected, the genes are converted to their human orthologs via the Ensembl 109 database (Feb. 2023)^65^.

**Figure 1:**
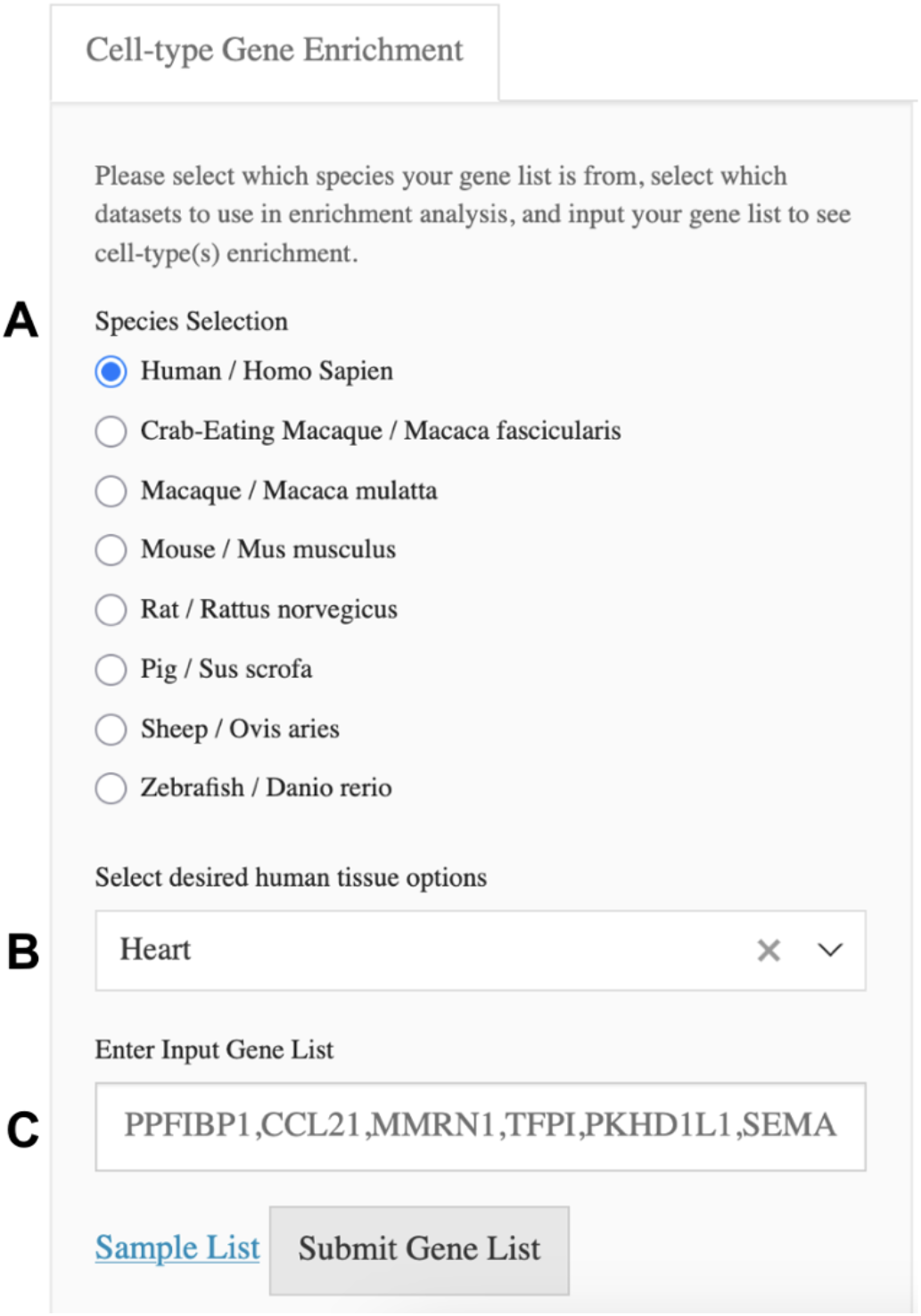
Inputs for celltypeEnrich. There are three inputs: **(A)** species of input gene list which determines if gene orthologs need to be found, **(B)** tissue(s) of interest which control which reference datasets are used in the analysis, and **(C)** input gene list. **Alt Text**: Screenshot of input side panel for web application. The three input selections are labeled A through C.

The second input section requires the user to select the tissue(s) to include in the enrichment analysis (Figure 1B). Currently, celltypeEnrich includes 26 reference datasets covering 43 tissues. The list of tissues is categorized by organ system, and users can select all tissues (default), all tissues within a system, individual tissues, or any combination thereof.

Finally, users must enter their genes in the provided input field, either one per line or comma-separated (Figure 1C). The gene list can be any combination of Ensembl IDs or gene symbols. While celltypeEnrich requires only five genes for enrichment analysis (after ortholog gene conversion), more genes are recommended to ensure unbiased results. A sample gene list of lymphatic endothelial markers from Litviňuková et.al can be used by clicking the words ‘Sample List.’ Clicking ‘Submit Gene List’ will then execute the enrichment analysis.

#### 3.1.2) Description of results

The enrichment results include five components (Figure 2). The top horizontal section (Figure 2A) has the list of input genes that were either not identified in the Ensembl database or did not have a human ortholog. If there is an error, such as not enough input genes or no significant results, the error message will be displayed in this section. The middle horizontal section (Figure 2B) allows the user to enter a file name if they want to download a CSV file of the significant enrichment results. The last horizontal section displays the main results in three tabs where each tab shows a different result format. The first tab (Figure 2C) displays enrichment plots for each dataset used in the enrichment analysis. The sidebar on the left of this tab shows all of the datasets with significant enrichment of at least one cell type. The enrichment plots’ x-axis is the negative log10-adjusted p-value, and the y-axis shows the four cell types with the most significant enrichment. The second tab (Figure 2D) shows a bar plot of the consensus cell-type annotation with the percent frequency each word occurs across all top results (see Methods 2.3). The bar plot includes a separate section for each hierarchical level of the cell-type annotation. The third tab (Supplemental Figure 1) is the full table of significant results (which can also be downloaded, see Figure 2B). It includes the negative log10-adjusted p-values, the cell types that are enriched, the dataset(s) that gave the significant enrichment, and the genes (in both Ensembl id and gene symbol formats) from the input list that were cell-type-specific. If ortholog conversion occurred, then the cell-type-specific genes reported are the human orthologs.

**Figure 2.**
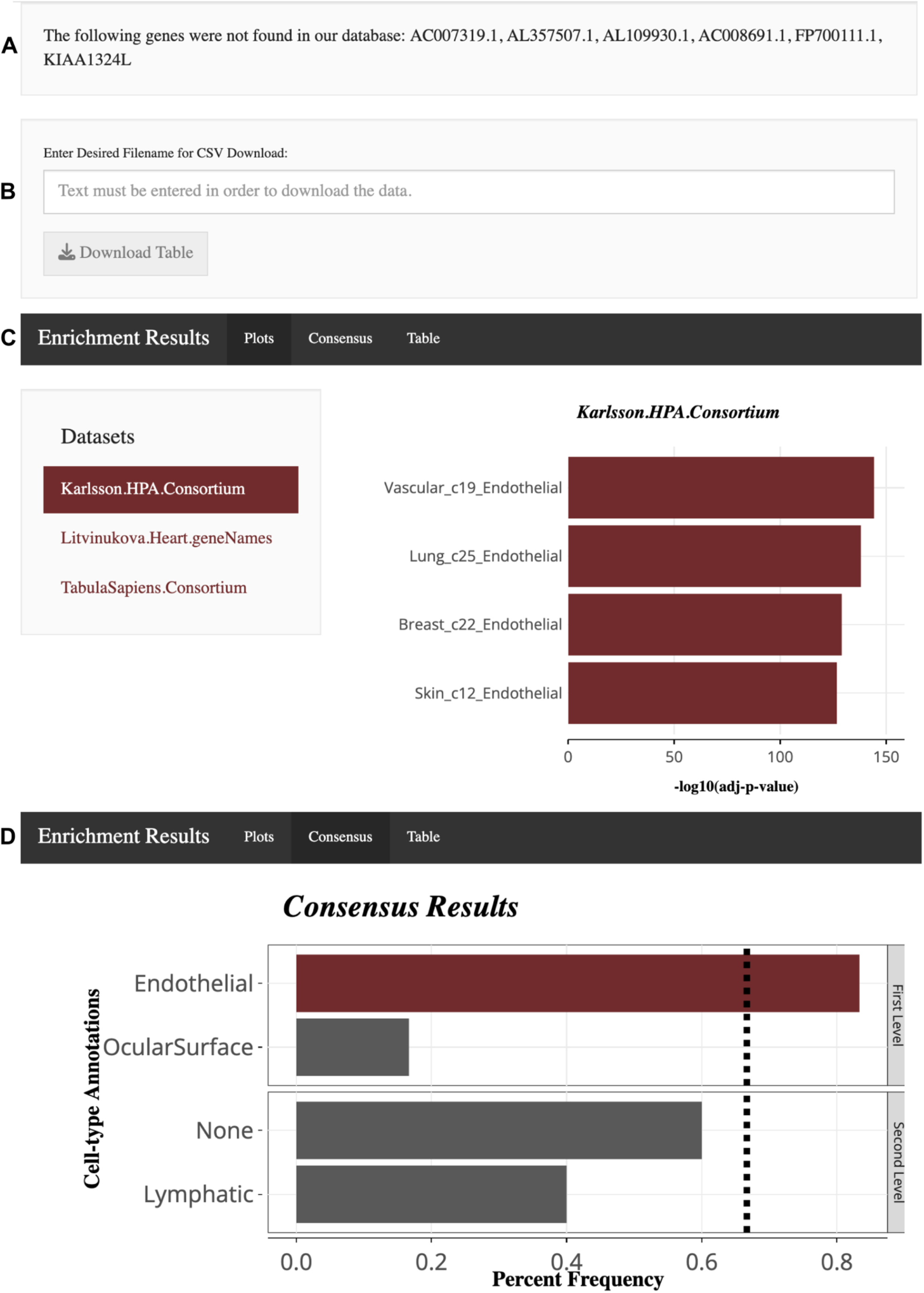
Example Results for Endothelial Lymphatic markers from Litviňuková et.al. The error message box **(A)** and the download CSV file box **(B)** are above the different results. An example set of results are shown in **(C)** and **(D).** There are three result output formats. Individual dataset enrichment plots which show the top 4 significantly enriched (highest -log10 adjusted p-values) cell types per reference dataset **(C)** and consensus results which determines if repeated enrichments have occurred across the reference datasets **(D)** are shown here. All enrichments are shown in table format in Supplemental Figure 1. **Alt Text**: Screenshots of example results output for web application. The four output boxes are labelled A through D.

### 3.2) Evaluating celltypeEnrich

#### 3.2.1) Benchmarking celltypeEnrich against existing methods

We benchmarked celltypeEnrich against other annotation tools using three scRNA-seq datasets (Adams Lung^66^, Koenig Heart^67^, and Scott Placenta^68^), none of which were used in developing celltypeEnrich. For both the lung and heart datasets, any disease-state cells were removed, and any cells labeled as unknown or as a multiplet were excluded from the comparison. Only cell types with at least 100 annotated cells were included in the comparison. Cluster markers were then found using the FindAllMarkers function in Seurat^36^. The placenta dataset only included control-state cells, and we therefore used the cluster markers provided in the original publication. Because the placenta dataset was from rat, which is not directly supported by other tools, the rat genes were converted to their human orthologs using the Ensembl 109 database^65^, and only genes that had orthologs were used for analysis in those tools. In total, 25 cell-types/cluster marker lists were used for benchmarking from the Adams lung dataset, 13 from the Koenig heart dataset, and 10 from the Scott placenta dataset, resulting in 48 total marker lists.

EasyCellType^3^ and SingleR^4^ each offer multiple options for databases or reference datasets to use for cluster annotation. We first evaluated the available reference databases/datasets within each tool and selected the best-performing option (Fisher’s Exact test with the Panglao database for EasyCellType; the He Organ Atlas and Zillonis Tumor Lung reference datasets for SingleR) for downstream benchmarking against celltypeEnrich (Supplemental Figure 2, See Methods 2.4).

When using the consensus results from celltypeEnrich (calculated using all reference datasets relevant to the test dataset), we found that it had relatively consistent accuracy (62-72%) in annotating marker lists, which was not the case for other tools that were tested (Figure 3).

**Figure 3:**
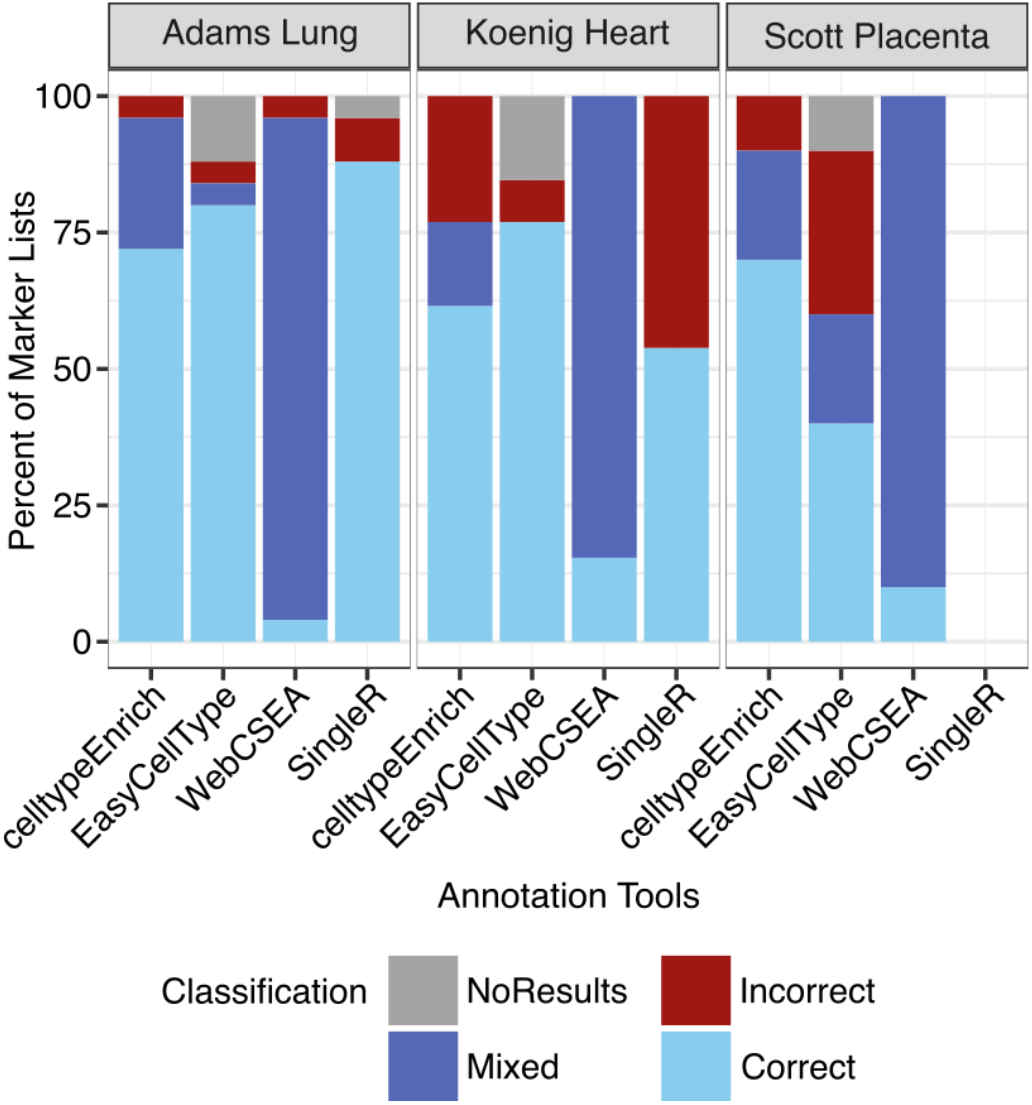
Accuracy of cell-type annotation between celltypeEnrich, EasyCellType, WebCSEA, and SingleR. The percent of Correct, Incorrect, Mixed, and No Results across all the annotated cell types’ differentially expressed genes from the Adams Lung, the Koenig Heart, and the Scott Placenta datasets. Tabulation of these results are in Supplemental Table 5. The classification of correct was based on published annotations, whereas Mixed results occurred when both Correct and Incorrect annotations were significant. **Alt Text**: A bar chart showing the percent of annotations classified as Correct, Incorrect, Mixed, or No Results from four tools: celltypeEnrich, EasyCellType, WebCSEA, and SingleR.

For the Adams lung dataset, SingleR correctly annotates four more cell types than celltypeEnrich, while EasyCellType correctly annotates two more cell types compared to celltypeEnrich (see Supplemental Table 5 for tabularization of Figure 3). However, SingleR incorrectly annotated two cell types, whereas both celltypeEnrich and EasyCellType incorrectly annotated one cell type. In addition, both SingleR and EasyCellType did not have annotations for one and three marker lists respectively, while celltypeEnrich always had an annotation. WebCSEA had mostly Mixed results (92%). For the Koenig heart dataset, celltypeEnrich correctly annotated one more cell type than SingleR. EasyCellType correctly annotated two more cell types than celltypeEnrich. Again, SingleR incorrectly annotates more cell types (6) than both celltypeEnrich (3) and EasyCellType (1). EasyCellType did not have annotations for two marker lists, while again celltypeEnrich always had an annotation. WebCSEA had 85% Mixed results. For the Scott placenta dataset, SingleR does not have a built-in placental model and was not included in the comparison. celltypeEnrich correctly annotated three more cell types than EasyCellType while also incorrectly annotating two less cell types. In addition, EasyCellType did not have annotations for one marker list, while again celltypeEnrich always had an annotation. WebCSEA again had mostly Mixed results (90%). Overall, SingleR incorrectly annotated twice as many marker lists as celltypeEnrich. SingleR’s accuracy was either 88% or 54% depending on the reference dataset used, whereas celltypeEnrich achieved a more consistent accuracy, ranging from 62% to 72% across benchmark datasets. While EasyCellType, had higher accuracy than celltypeEnrich on two of the three benchmark datasets when using its highest-performing database and statistical test combination, when using its default database and statistical test combination (CellMarker database and GSEA test), EasyCellType had drastically lower accuracy, ranging from 0% to 16% (Supplemental Figure 2). These results suggest that EasyCellType’s performance is dependent on database selection, whereas celltypeEnrich provides more consistent performance without requiring optimization.

#### 3.2.2) Robustness of celltypeEnrich to downsampling

We performed a robustness test for celltypeEnrich to determine if performance is a>ected as the number of input genes decreases. Each of the marker gene lists for each of the three datasets used in the benchmark comparison was randomly sampled 100 times to 75%, 100 times to 50%, and 100 times to 25% of its original gene list size. Since the original marker gene lists ranged in size from 115 to 3043 genes, the 75% downsampled marker lists ranged in size from 87 to 2283 genes, the 50% downsampled marker lists ranged from 58 to 1522 genes, and the 25% downsampled marker lists ranged from 29 to 761 genes (Figure 4A). All gene lists were analyzed using celltypeEnrich, and performance was evaluated as described in the last paragraph of Methods 2.2.

**Figure 4:**
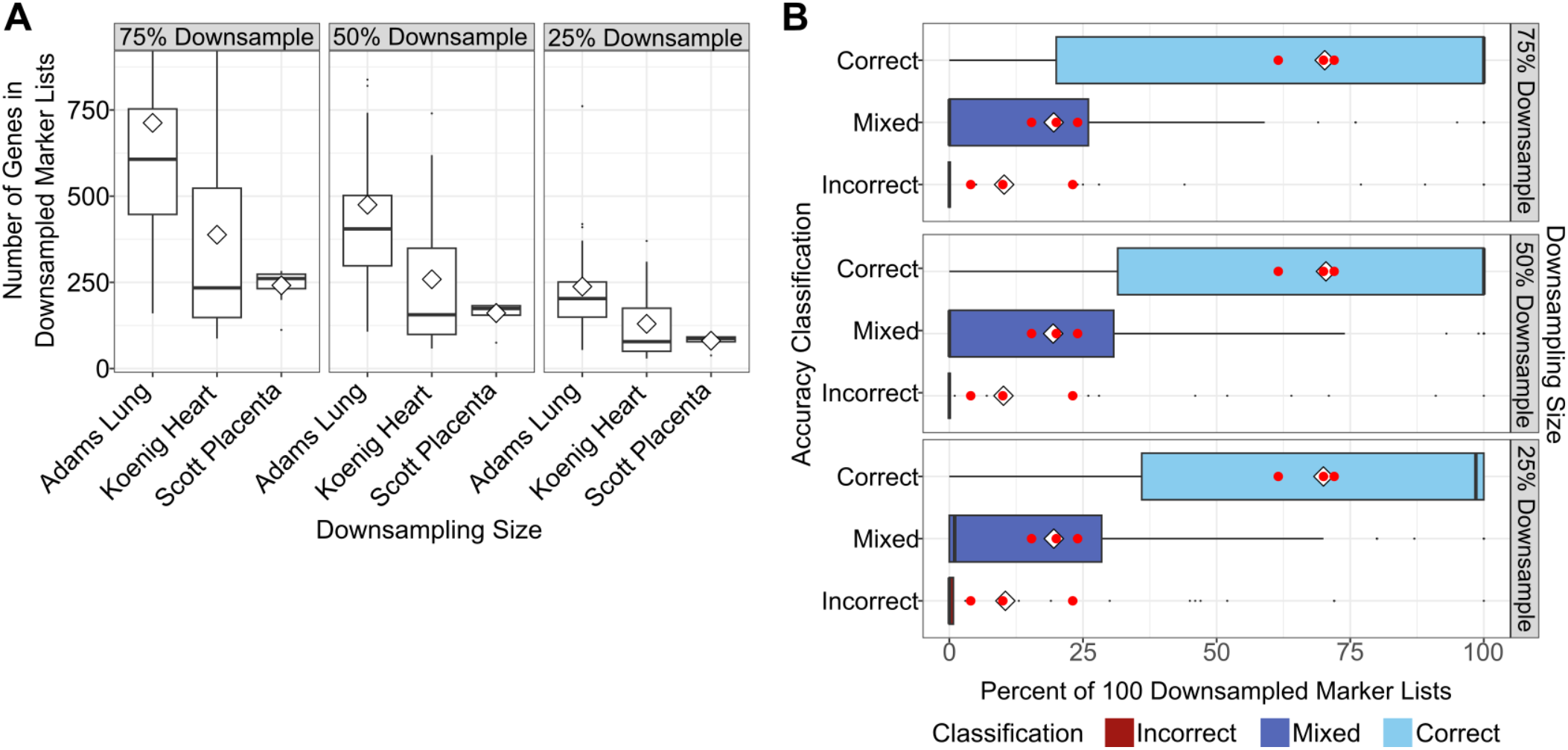
Determining robustness of celltypeEnrich. **(A)** Number of genes in each downsampled marker list for each dataset. Separate plots are shown for marker lists downsampled to 75%, 50%, and 25% of their original size. The white diamond is the average number of genes across marker lists in the dataset. **(B)** celltypeEnrich performance across 100 randomly downsampled marker gene lists per cell type. Marker lists were downsampled to 75%, 50%, or 25% of their original size, with each downsampling level shown in separate plots. The percentage of Correct, Mixed, and Incorrect annotations is plotted for downsample. The white diamonds are the average percent of Correct, Mixed, and Incorrect annotations across all cell type marker lists and all downsamples. The red dots are the corresponding percentages obtained using the original marker lists (Figure 3) for each dataset. The classification of Correct was based on published annotations, whereas Mixed results occurred when both Correct and Incorrect annotations were significant. **Alt Text**: Graphs showing (A) size and (B) percent of Correct, Mixed, and Incorrect results for down-sampling of marker lists.

For each marker gene list across all datasets, the percentage of downsampling iterations classified as Correct, Mixed, or Incorrect was calculated (Figure 4B). Across the three downsampling levels, the median percentage for Correct results varied by less than 1.5%, and the average percentage varied by less than 1%. The median percentage of Correct annotations was 100% for the 75% and 50% downsampling levels and decreased slightly to 98.5% for the 25% downsampling level. Similarly, the average percent of Correct annotations remained stable at 70-70.5% for all three downsampling levels. Although the median and average percentages changed minimally as the marker lists were downsampled, the interquartile range of Correct annotations narrowed with increasing downsampling. For the 75% downsampling level, the interquartile range for Correct annotations spanned from 20% to 100%, compared with 31.5% to 100% for the 50% downsampling level and 36%-100% for the 25% downsampling level.

Given how broad the interquartile range of Correct results per cell type was, we investigated these results by cell type (Figure 5). 30 out of the 48 total marker lists (across all cell types for the 3 datasets used for benchmarking) had Correct annotations 90-100% of the time across all three down-sampling sizes. Surprisingly, only two marker lists (Macrophage from the Adams lung dataset and SmoothMuscle from the Scott placenta dataset) had more than a 10% decrease in the percent of Correct annotations as the marker list size decreased. More in line with the observed interquartile range changes from Figure 4B, six marker lists (Alveolar Macrophage and Cytotoxic T cells from the Adams lung dataset, Epicardium and Pericytes from the Koenig heart dataset, and Macrophage clusters 17 and 18 from the Scott placenta dataset) had more than a 5% increase in the number of Correct annotations at either the 50% or 25% downsampling levels. Overall, relatively few marker lists showed marked declines in performance with downsampling, supporting the robustness of celltypeEnrich consensus annotations.

**Figure 5:**
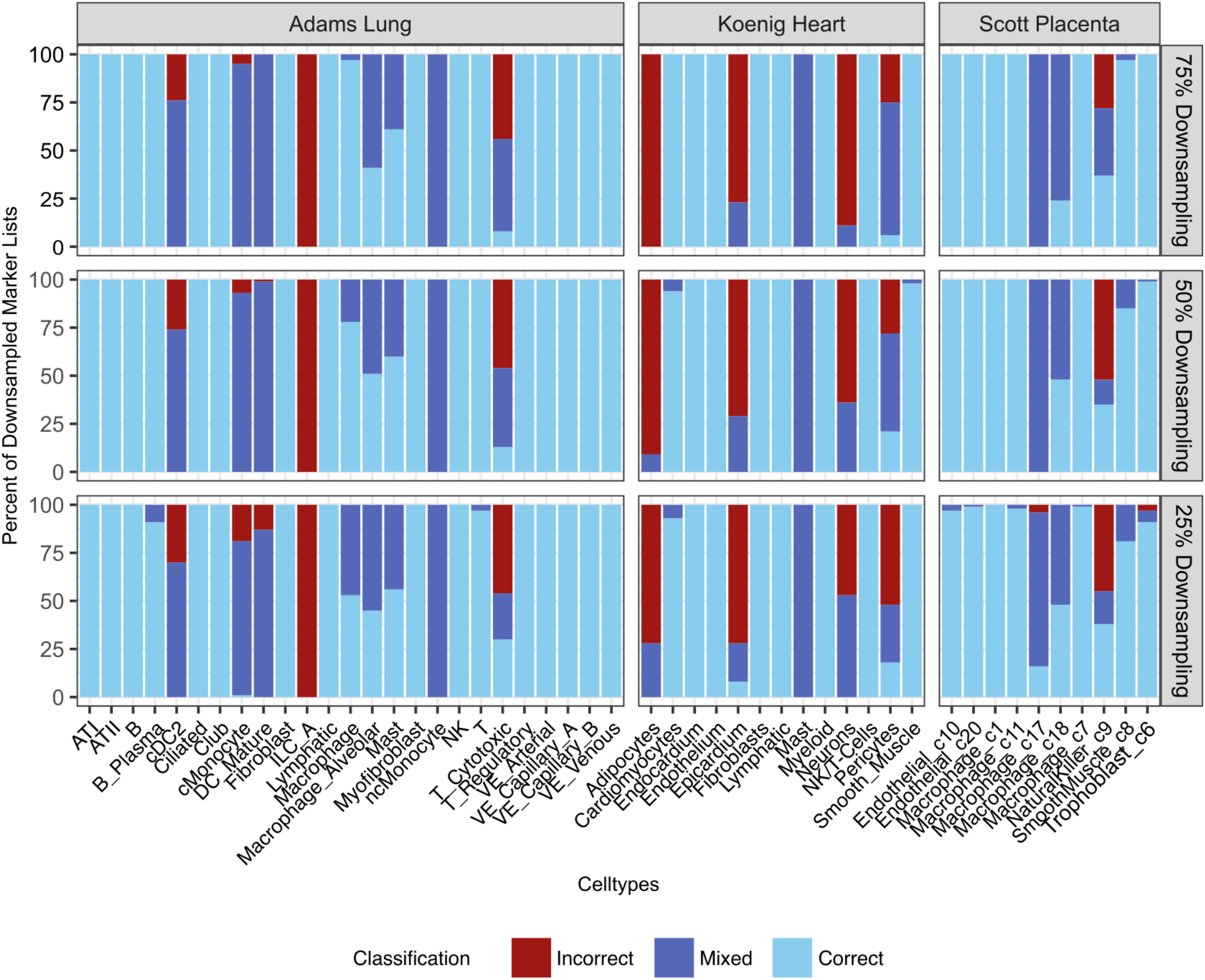
celltypeEnrich’s performance across downsampled marker lists by cell type and dataset. Cell types are listed in alphabetical order by dataset. Separate plots are shown for marker lists downsampled to 75%, 50%, and 25% of their original size. The classification of Correct was based on published annotations, whereas Mixed results occurred when both Correct and Incorrect annotations were significant. **Alt Text**: Bar chart showing percent of Correct, Mixed, and Incorrect results for each cell type’s down-sampled marker lists.

## 4) Discussion

celltypeEnrich is a cluster-level annotation tool that determines whether an input marker gene list is significantly enriched for cell-type-specific gene sets determined from multiple publicly available datasets. It then calculates a consensus annotation based on results across each cell-type-specific gene set analyzed. By using repeated enrichment signals across datasets, celltypeEnrich reduces reference-dataset-specific bias and increases confidence in the assigned cell-type labels. Using three independent scRNA-seq datasets, we benchmarked celltypeEnrich against EasyCellType, WebCSEA, and SingleR and found that celltypeEnrich was more consistent in correctly annotating than these tools. While EasyCellType correctly annotated 2 more cell types in two of the benchmark datasets, this is contrasted with both 40% accuracy for the third dataset and a less than 20% accuracy when using the default reference database. celltypeEnrich offers reliable annotation no matter the tissue of interest and without relying on specific parameter combinations. While SingleR had comparable performance with one dataset, celltypeEnrich offers greater flexibility by not relying on tissue-specific models or being restricted to human and mouse input data. Instead, celltypeEnrich can incorporate enrichment data from 43 tissues and supports ortholog conversion for seven non-human species, making it broadly applicable across tissues and when understanding how data from multiple model organisms relates to human cell types. Together, the consensus-based results and broad reference coverage distinguish celltypeEnrich from many existing annotation tools.

When relabeling cell types across datasets, we observed that the consortium datasets typically provide broader cell-type annotations, whereas focused single-tissue scRNA-seq studies often include more specialized cell-type subtype annotations. Although cell-type labels in celltypeEnrich are derived from published annotations, which are often based on canonical marker genes, the use of multiple independent datasets with varying annotation specialization mitigates reliance on any single annotation scheme. By using both the consortium and single-tissue datasets in the enrichment analyses, celltypeEnrich benefits from the complementary strengths of each: the broad tissue-wide coverage of consortium datasets and the finer cell-type resolution from single-tissue studies. Analyzing enrichment results across these datasets further reduces the possibility of misannotation by requiring consistency from multiple independent sources.

Relatedly, in contrast to manual annotation approaches, which often rely on a small number of canonical marker genes to label clusters, celltypeEnrich calculates enrichment using large gene sets. By using expression information across many genes, the influence of overlapping expression of individual marker genes from different cell types is reduced, decreasing the likelihood of incorrect cluster annotations. Together, these features enable celltypeEnrich to provide more robust and biologically consistent annotations across diverse datasets.

To assess the robustness of celltypeEnrich, we evaluated performance under systematic downsampling of input marker gene lists and observed minimal overall variation in annotation outcomes as marker list size decreases. However, cell-type-specific trends emerged when examining individual annotations (Figure 5). Myeloid immune cell types were more likely to have Mixed annotations at least 50% of the time across all downsampled sets. Tissue-specific or rare cell types (i.e. ones that might not be present in multiple datasets, such as Innate Lymphoid Cells in the lung dataset) had Incorrect annotations at least 50% of the time across all downsampled sets. Interestingly, in some cases at the 75% downsampling level, cell types had mostly Mixed annotations, whereas at the 25% downsampling level the same cell type had more Correct and Incorrect annotations. This could be the result of downsampling an initially mixed list of genes, where there were cell-type-specific genes for both the correct cell type and at least one incorrect cell type. When down-sampling to 75% of the original size, the possibility of sampling cell-type-specific genes for only the correct cell type is low because the number of genes being excluded is low. When downsampling to 25% of the original size, a smaller number of genes is selected, increasing the possibility of sampling just the cell-type-specific genes for the correct cell type. The same logic can be applied to the cell-type-specific genes of the incorrect cell type(s). As a result, what was initially a Mixed classification could result in either Correct or Incorrect classifications.

Immune cell types accounted for the majority of cases with Mixed or Incorrect annotations across downsampling levels, suggesting that a substantial fraction of cell-type-specific genes are shared across immune lineages in different datasets. In celltypeEnrich, cell-type classification depends not only on annotation accuracy but also on the cell types defined in the reference dataset. While it is difficult to determine how accurate published cell-type annotations are, previously published work has shown that distinguishing between T cell subtypes is particularly challenging^69^. This might suggest that broad cell-type annotations are more reliable than the more specific subtype annotations that distinguish between closely related cell-type populations. In the benchmarking and robustness analyses, any subtype matching the expected annotation was considered a Correct annotation (i.e. Memory CD4, Regulatory CD4, Activated CD8, and GD T cell annotation results would be considered Correct for a T cell marker list). However, this does not address the issue of cell-type-specific genes being shared between different cell types, such as Monocytes and Neutrophils.

Busarello et al. have previously highlighted the challenge that different datasets annotate cell types at varying levels of granularity^70^, and we observe this issue in the reference datasets included in celltypeEnrich. For example, in the endometrium dataset^19^, immune cells were annotated only at the broad level of myeloid and lymphoid lineages, whereas the gut dataset^17^ included 19 immune cell types, but lacked annotations for certain immune populations such as natural killer cells. Such inconsistencies in annotation level can lead to increased Mixed or Incorrect annotations. In the case of natural killer cells, for instance, they may be labeled as lymphoid cells or misclassified as either B cells or T cells, depending on the reference datasets used, contributing to ambiguity in enrichment-based annotations.

In addition to annotation inconsistencies, biological similarity among immune cell types likely contributes to observed overlap in cell-type-specific gene sets. For example, natural killer T cells are expected to share transcriptional features with both natural killer cells and T cells, complicating precise discrimination. These challenges reflect fundamental limitations in current reference annotations and underlying biological complexity, rather than shortcomings of celltypeEnrich itself. Addressing such issues will require improved consistency in cell-type definitions, expanded reference datasets, and coordinated meta-analyses across studies, which represent important directions for future work.

## 5) Conclusion

celltypeEnrich is a cluster-level annotation tool that uses the hypergeometric test to determine whether an input gene list is enriched for cell-type-specific genes. Its use of multiple tissue datasets enables high-confidence consensus annotations, distinguishing celltypeEnrich from other annotation tools. We have shown that celltypeEnrich provides robust, unbiased annotations and performs comparably or better than available annotation tools. As new work is published, celltypeEnrich can be readily updated to include additional datasets, including spatial transcriptomics data, which would provide additional biological context and further improve annotation confidence.

## 6) Acknowledgements

The authors thank the Tuteja Lab members for testing celltypeEnrich and reviewing the manuscript, as well as Miles Aronnax, the LAS research IT team at ISU, Misty Treanor, and the CALS/LAS Web team at ISU, for setting up the web application server.

## 8) Data Availability

All datasets used both in creating and testing celltypeEnrich were derived from sources in the public domain (See Supplementary Table 1 ‘Data Source URL’ Column). The celltypeEnrich RShiny tool can be accessed at https://celltypeenrich.gdcb.iastate.edu. The RShiny code is available at https://github.com/Tuteja-Lab/celltypeEnrich.

