## Supplementary Figures for "celltypeEnrich: a consensus-based scRNA-seq cluster annotation tool"

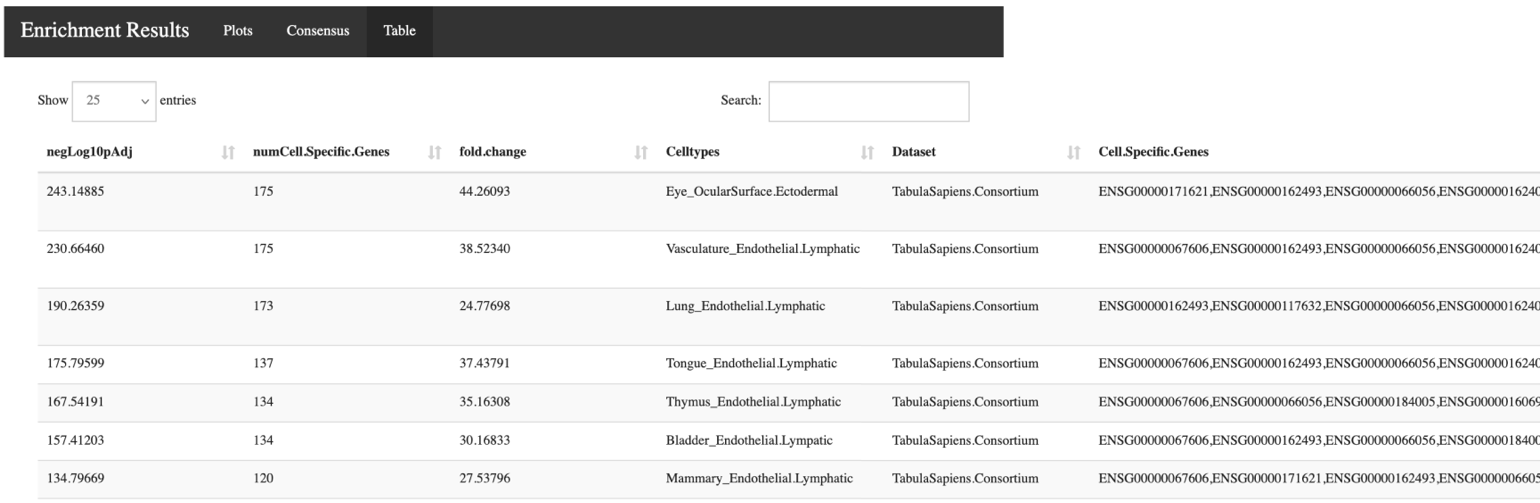

**Supplemental Figure 1**: Example Screenshot of Table Results for Endothelial Lymphatic markers from Litviňuková et.al. The table shows the negative-log10-transformed adjusted p-value (negLog10pAdj), which indicates significance of enrichment for the cell type, the number of cell-type-specific genes from the input list for the cell type (numCell.Specific.Genes), the fold-change, the cell type, the reference dataset that the cell type originates from, and the cell-type-specific genes leading to the cell-type enrichment (this is cut off due to the size of the website).

**Alt Text:** Screenshot of example table results.

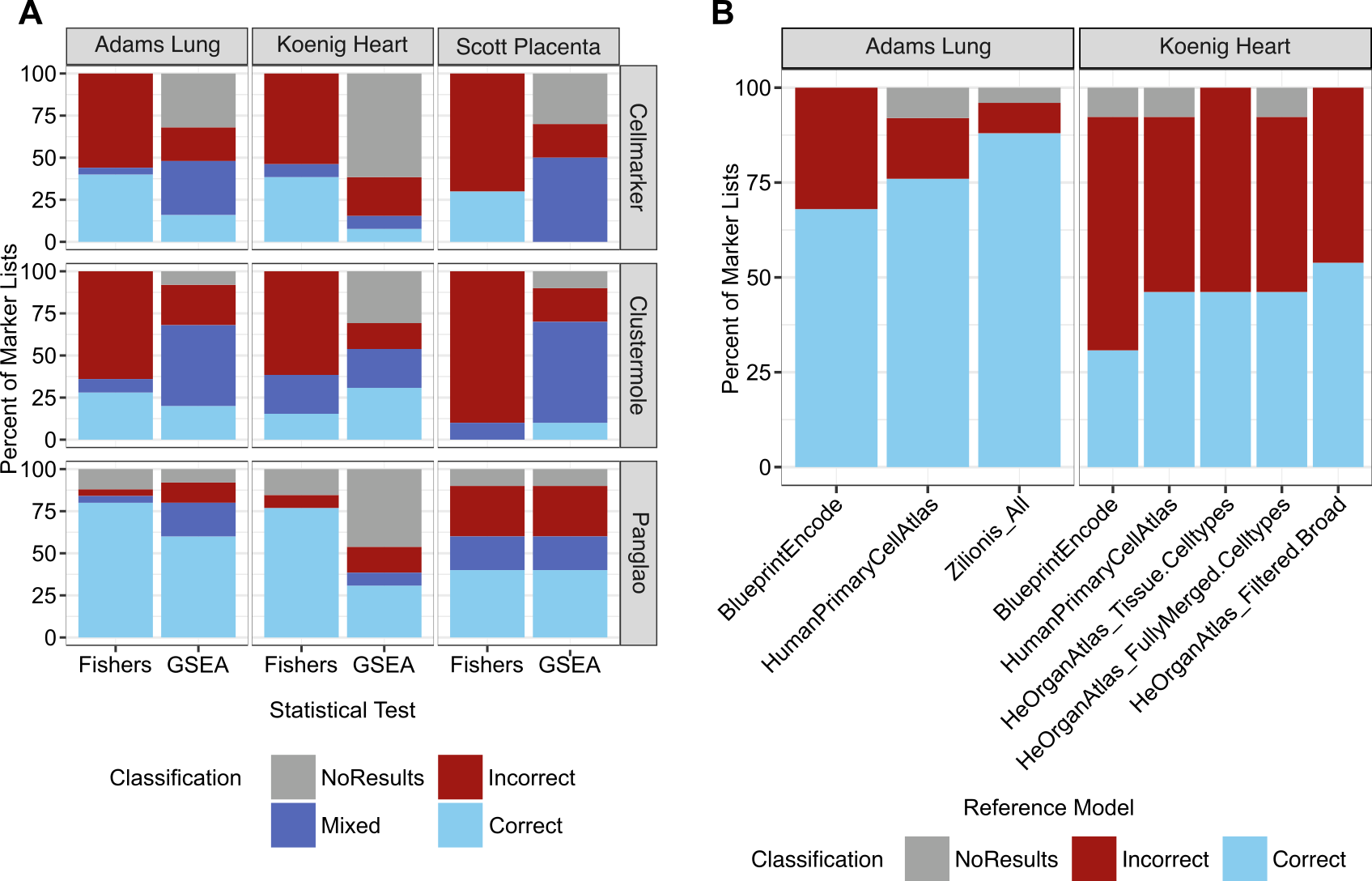

**Supplemental Figure 2**: Evaluating input parameters for Easy Cell Type and SingleR. For **(A)** and **(B)** the percent of correct annotations based on published annotations was used to determine optimal parameter combination. Mixed results occurred when both Correct and Incorrect results were significant. **(A)** Modulating EasyCellType’s parameters to determine which combination of statistical test (Fisher’s Exact or GSEA) and database (CellMarker, Clustermole, or Panglao) provides the best performance given the known annotations. Using the Panglao database with the Fisher’s Exact test gave the best results for all three benchmarking datasets. **(B)** Using different reference datasets provided by SingleR to determine the optimal annotation set. Using the Zilionis and He Organ Atlas gave the best results out of the relevant SingleR reference datasets.

**Alt Text**: Graphs comparing results with different parameter combinations or different reference datasets for EasyCellType and SingleR.
